# Development of an in vitro senescent hepatic cell model for metabolic studies in aging

**DOI:** 10.1101/2020.03.31.014035

**Authors:** Brijesh Kumar Singh, Madhulika Tripathi, Reddemma Sandireddy, Keziah Tikno, Jin Zhou, Paul Michael Yen

**Author notes:** Contact Cardiovascular and Metabolic Disorders Program, Duke-NUS Medical School, Singapore 169857.

## Abstract

Although aging in the liver contributes to the development of chronic liver diseases such as NAFLD and insulin resistance, little known about the molecular and metabolic details of aging in hepatic cells. To examine these issues, we used sequential oxidative stress with hydrogen peroxide to induce premature senescence in AML12 hepatic cells. The senescent cells exhibited molecular and metabolic signatures, increased SA-βGal and γH2A.X staining, and elevated senescence and pro-inflammatory gene expression that resembled livers from aged mice. Metabolic phenotyping showed fuel switching towards glycolysis and mitochondrial glutamine oxidation as well as impaired energy production. The senescent AML12 cells also had increased mTOR signaling and decreased autophagy which likely contributed to the fuel switching from β-oxidation that occurred in normal AML12 cells. Additionally, senescence activated secretory proteins from conditioned media of senescent cells sensitized normal AML12 cells to palmitate-induced toxicity, a known pathological effect of hepatic aging. In summary, we have generated senescent AML12 cells which displayed the molecular hallmarks of aging, and also exhibited the aberrant metabolic phenotype, mitochondrial function, and cell signaling that occur in the aged liver.

## INTRODUCTION

Aging is a major risk factor for many chronic diseases. In the liver, aging increases the susceptibility towards acute liver injury and hepatic fibrotic response [1–3]. Moreover, aging has been positively associated with increased risk and poor prognosis of various liver diseases including non-alcoholic fatty liver disease (NAFLD), insulin resistance, alcoholic liver disease, hepatitis C, and negatively associated with hepatic regenerative capacity [3, 4]. Currently, the study of aging and chronic hepatic diseases has been hampered by the long period of time necessary to conduct human and animal studies, and the limited relevance of non-mammalian models to human diseases. While there are *in vitro* aging models that employ fibroblasts, there currently are no reliable *in vitro* hepatic cell models to study aging in the liver.

Cellular senescence in the main feature manifested in tissues of the aging organism [1, 5–8]. It is characterized by permanent cell cycle arrest, resistance to apoptosis, and a senescence-associated secretory phenotype [5]. Under pathological stress conditions, excessive accumulation of senescent cells in affected tissues adversely affects their regenerative ability and creates a pro-inflammatory environment that can resemble those found in age-related disorders such as Alzheimer’s disease, cardiovascular disease, Type 2 diabetes, and other conditions including chronic liver diseases [3, 5, 7, 9–12]. Targeting senescent cells has the potential to delay age-associated disorders and/or reverse pathological metabolic phenotypes [3, 13–16]. Recent studies in the liver show that inducing hepatocyte senescence promotes fat accumulation and hepatic steatosis *in vitro* and *in vivo* [17]. Likewise, targeting senescent hepatocytes and adipocytes reduces overall hepatic steatosis and improves obesity-induced metabolic dysfunction [14, 17]. These findings suggest that senescence plays an important role in the development of chronic hepatic diseases and their metabolic abnormalities.

Previous senescent cell models have used primary fibroblast or hepatic cancer cells that were subjected to oxidative stress [18–22], or primary hepatocytes that underwent gamma irradiation [17]. However, these models have certain limitations since different cell types will react differently to a given stressor [23]. Furthermore, the method of senescence induction *i.e.,* oxidative stress, gamma irradiation or overexpression of an oncogene, may contribute to variable phenotypes that may not necessarily resemble aging *in vivo* [5, 16, 23]. Thus, it is critical to compare and verify the fidelity of *in vitro* models with tissues from aged mice. Although there still remains some uncertainty about the optimal method to induce senescence *in vitro*; it generally is agreed that oxidative stress and mitochondrial dysfunction play significant roles in the aging process [6, 11, 24, 25]. Using repetitive H_2_O_2_ exposures, we generated hepatic cellular senescence in hepatic Alpha Mouse Liver 12 (AML12) cells. Our characterization of the senescent cells showed that these senescent AML12 cells exhibited age-related molecular and metabolic changes, particularly a fuel switch to glycolysis and glutamate oxidation and decreased autophagy found in the livers of aged mice.

## RESULTS

### Senescence induction and validation in AML12 cells

AML12 cells were grown to 50% confluency and then treated with 1 mM H_2_O_2_ for 1 h for one day. They subsequently were treated with 750 μM H_2_O_2_ in serum-free DMEM:F12 medium for 1 h per day for 5 consecutive days as described in **Figure 1a**. After each treatment, serum-free DMEM:F12 medium containing H_2_O_2_ was replaced with complete DMEM:F12 medium (containing 10% FBS, 1x ITS, 100 nM dexamethasone, and 1x penicillin and streptomycin) for a 23 h period of recovery during each day. Cells then were sub-cultured/re-seeded in the ratio of 1:3 until they reached confluency >80% after approximately 3 days. As shown in **Figure 1b**, morphological effects such as cellular hypertrophy were after 2 days of H_2_O_2_ treatment. After 6 days of multiple H_2_O_2_ treatments, the cells showed classical features of senescence as they were distinctly larger and showed less proliferation than untreated controls (**Figure 1b,c**) [8]. Moreover, the mRNA expression of senescence marker genes, *TP53/p53*, *CDKN2A*/*p21* and *CDKN1A*/*p16* were increased in the senescent AML12 cells compared to control AML12 cells (**Figure 1d**). Similarly, the expression of these genes also was increased in livers from old mice (100-108 week age) compared to those from young mice (12-20 week age) (**Figure 1e**). Furthermore, multiple H_2_O_2_ treatments increased senescence since there was increased activated β-Gal (SA β-Gal)-positive cells (**Figure 2a,b**), γH2A.X-positive cells containing condensed chromatin in larger nuclei along with cellular hypertrophy (**Figure 2c-f**). Taken together, these data showed that multiple H_2_O_2_ treatments markedly induced premature senescence in AML12 cells [1]. Consistent to these findings, RNAseq analysis also showed that cellular senescence (P=0.009) and aging (P=0.012) pathways were significantly upregulated in senescent AML12 cells (Supplementary figure 1a).

**Figure 1.**
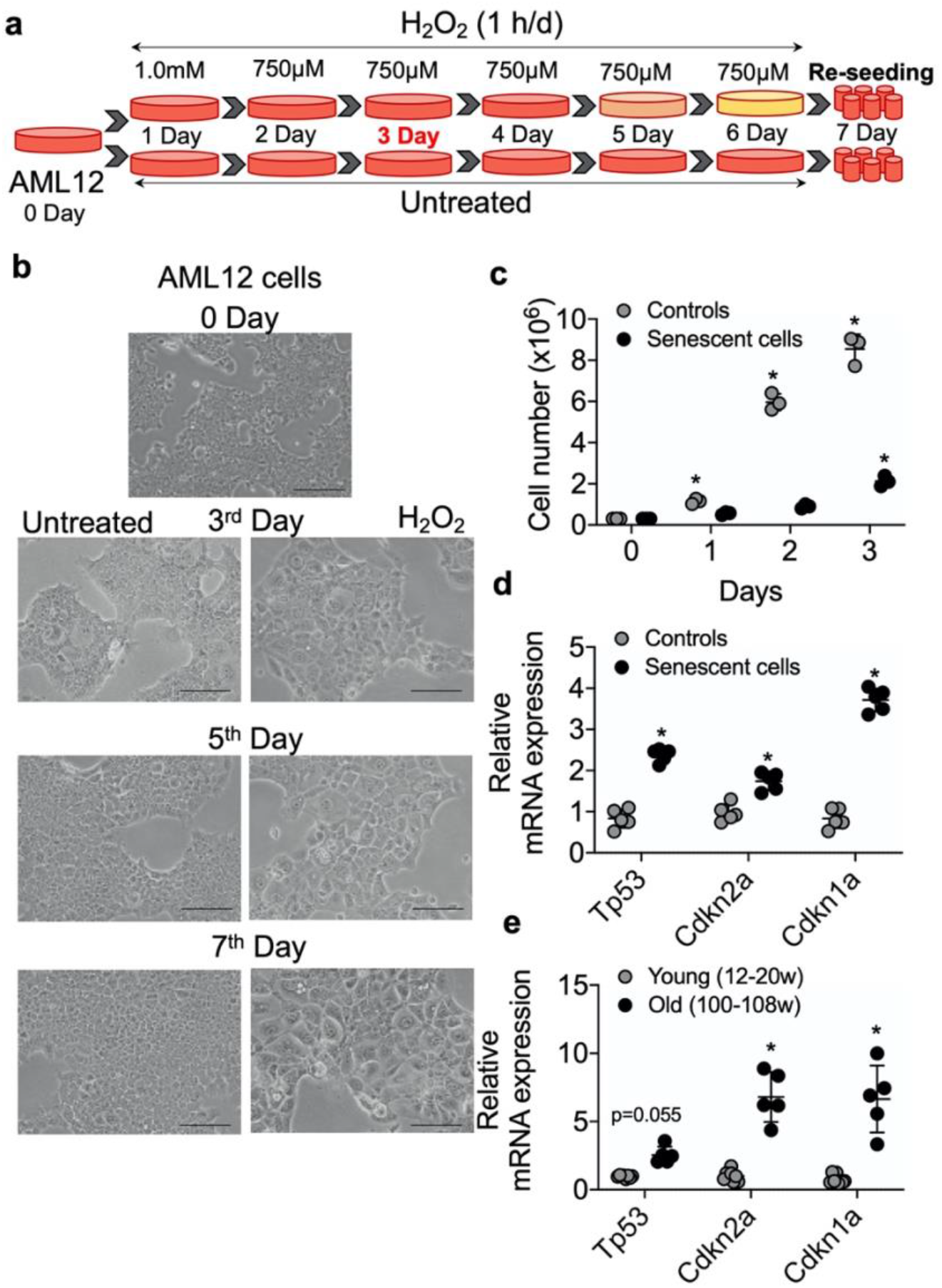
Senescence induction in mice normal hepatic cells AML12. (**a)** Schematic representation of experimental strategy for the senescence induction. 4×10_6_ Cells were seeded at day 0 in T175 cell culture flask. Next day (day 1) cells were treated with 1.0 mM H_2_O_2_ followed by 750 μM for the subsequent 5 days. Cells can be sub-cultured at 1:3 ratio at day 3 if required. At day 7, cells were re-seeded for experiments as required. (**b**) Visualization of senescence induction in mice AML12 cells during H_2_O_2_ treatments from day 3, day 5 and day 7 before re-seeding. The images are taken randomly at 10x magnification. Scale bars as 100 μm. (**c**) 0.3×10_6_ control and senescent AML12 cells were seeded in each well of the 6-well plate. Cells were trypsinized at indicated days and counted by an automated cell counter. (**d and e**) RT-qPCR analysis of senescence genes in AML12 cells (d) and liver tissues from young and old mice (e). * Statistical differences were calculated significant as p<0.05.

**Figure 2.**
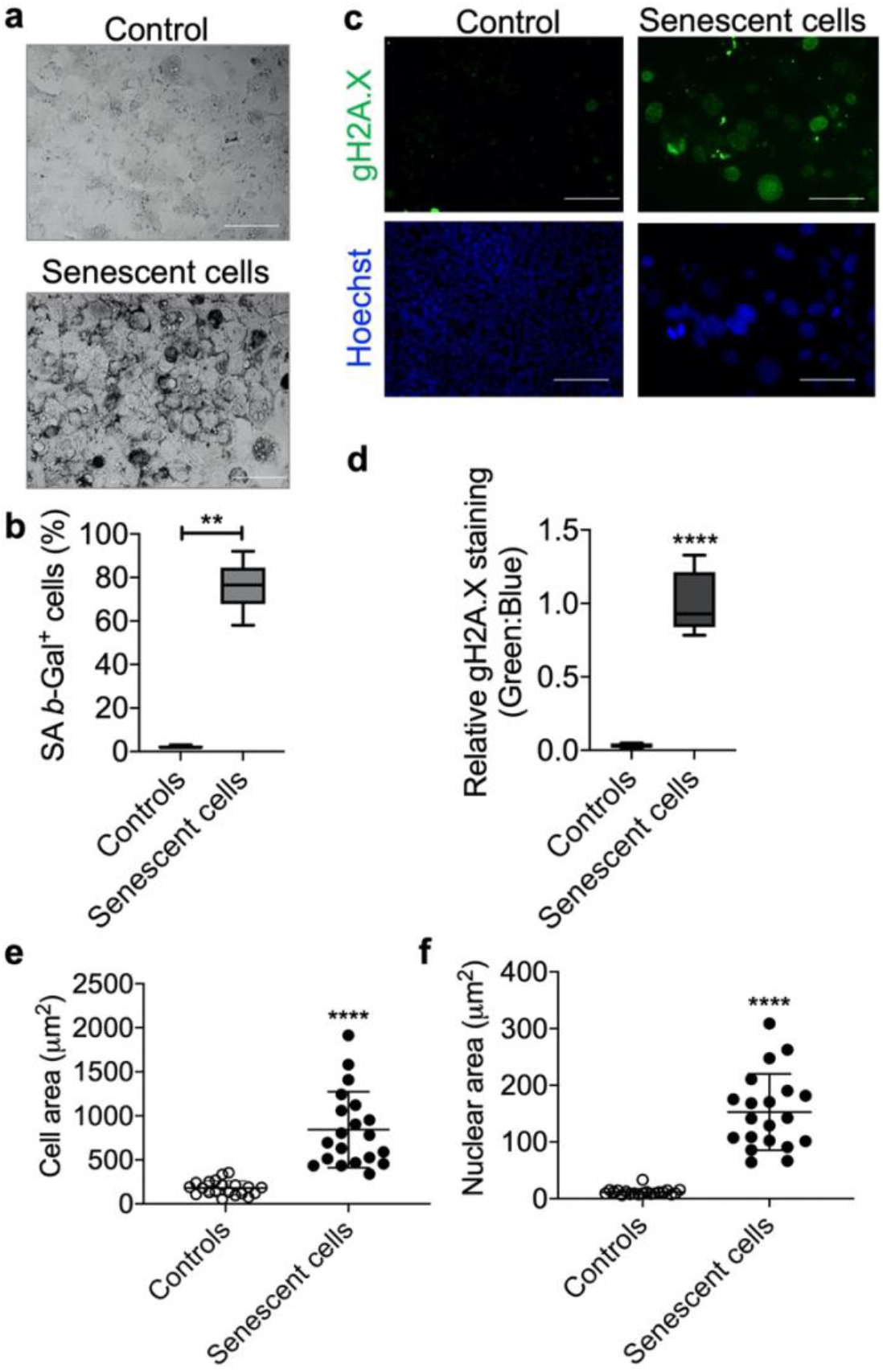
Senescence validation by SA β-Gal and γH2A.X staining. After senescence induction at day 7, cells were re-seeded in 4-chambered slides and next day SA β-Gal (**a**) and γH2A.X (**b**) staining were performed. Cells were counted manually for analysing SA β-Gal_+_ cells over total cells and presented as percent positive cells. Scale bars as 50 μm. (**c**) Cells were counter-stained with Hoechst 33324 for immunofluorescence imaging and relative γH2A.X signals were analysed over Hoechst 33324 signals. Scale bars as 100 μm. (**d**). (**e** and **f**). Area of the cells (e) and nucleus (f) were measured using ImageJ (NIH) software. Statistical differences were calculated significant as *p<0.05 and **p<0.001.

### Bioenergetic phenotype of senescent AML12 cells

To perform bioenergetic phenotyping of senescent AML12 cells, we used the Seahorse extracellular flux analyzer to measure the cell’s glycolytic and oxidative potential. Senescent AML12 cells showed higher oxygen consumption rate (OCR) and extracellular acidification rate (ECAR) than control cells (**Figure 3a**). Senescent AML12 cells also were more reliant on glycolysis for cellular energy than oxidative metabolism under energy stress conditions (**Figure 3b**). We further analysed the glycolytic potential of senescent AML12 cells and found that they had significantly higher basal glycolysis, glycolytic potential as well as glycolytic reserve capacity than control cells (**Figure 3c,d**). Furthermore, senescent AML12 cells had marked increases in basal mitochondrial activity and maximum respiratory capacity, but no change in mitochondrial ATP production (**Figure 4a,b**). Instead, these cells showed reduced coupling efficiency and increased proton leak, suggesting they had mitochondrial dysfunction, a hallmark of cellular senescence and aging. Finally, mitochondrial fuel oxidation analysis showed there were significant reductions in glucose and fatty acid flexibilities and capacities. In contrast, there was an enhancement of glutatmine oxidation flexibility and capacity (**Figure 4a-c)**. Similarly, glutamine-associated oxidation pathways were also upregulated in RNAseq analysis (Supplementary figure 1a, Supplementary data-RNAseq Pathway analysis> Pathways Up). However, there were no significant changes in glucose, fatty acid, or glutamine oxidation dependency (**Figure 4c**).

**Figure 3.**
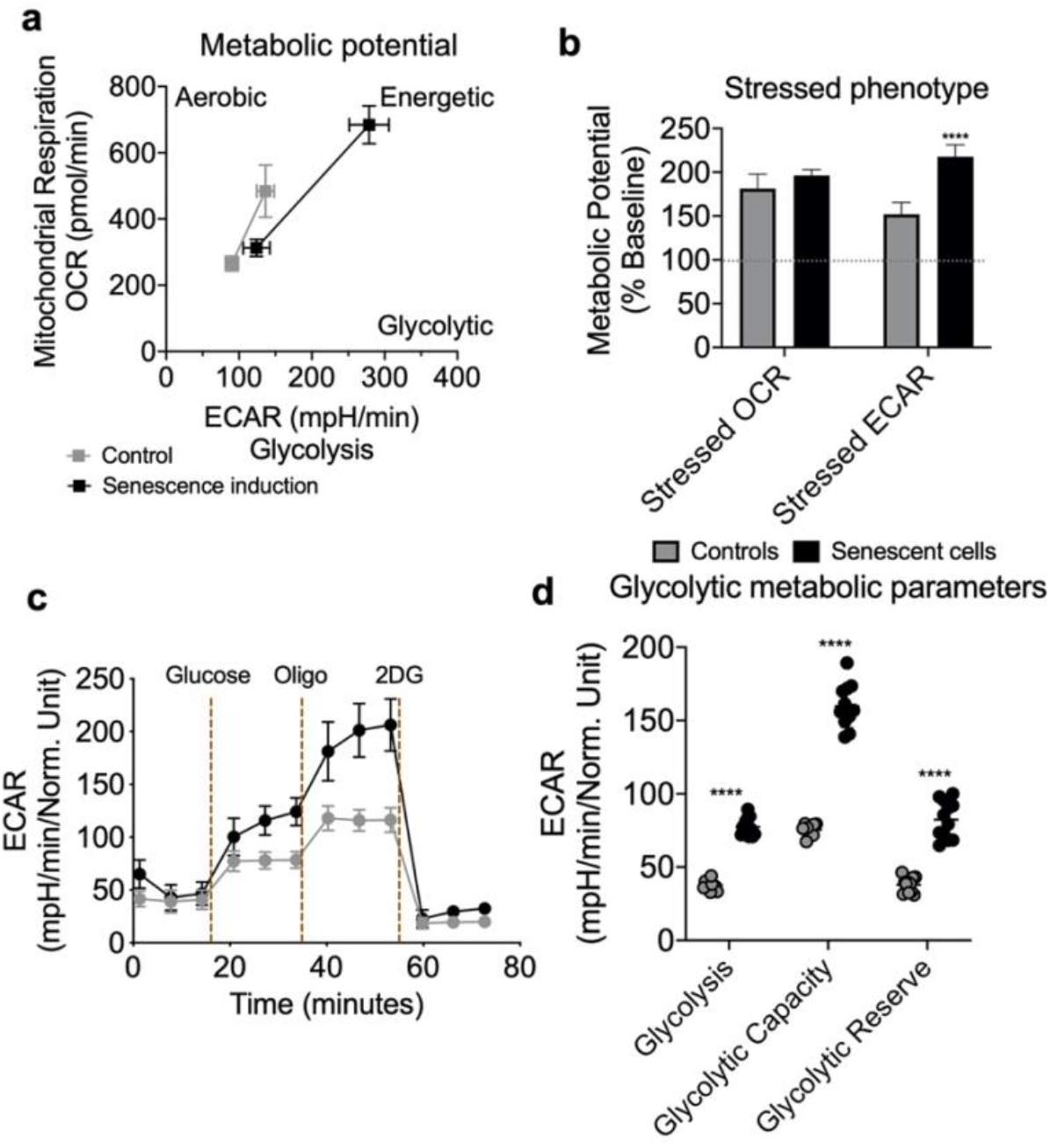
Seahorse extracellular flux analysis for metabolic potential and glycolytic flux in senescent AML12 cells. Agilent Seahorse XF Cell Energy Phenotype Test kit was used with Agilent Seahorse XFe96 Extracellular Flux Analyzer to analyse control and senescent AML12 cell’s metabolic potential (**a**) and stressed phenotype (**b**). Oxygen consumption rate (OCR) represents mitochondrial respiration while the extracellular acidification rate (ECAR) represents glycolytic potential under basal and stressed conditions (as described in Methods) (**a**). Percent change in mitochondrial oxidative phenotype (OCR) and glycolytic phenotype (ECAR) under stress over basal (100%) conditions (**b**). Agilent Seahorse XF Glycolysis Stress Test kit was used with Agilent Seahorse XFe96 Extracellular Flux Analyzer to analyse control and senescent AML12 cell’s glycolytic flux and reserve capacity (**c**). Glycolytic metabolic parameters were calculated as described in the Methods (**d**). All the parameters presented in the panel b and d were calculated using Seahorse Wave Desktop software. Statistical differences were calculated significant as **p<0.01; ****p<0.0001.

**Figure 4.**
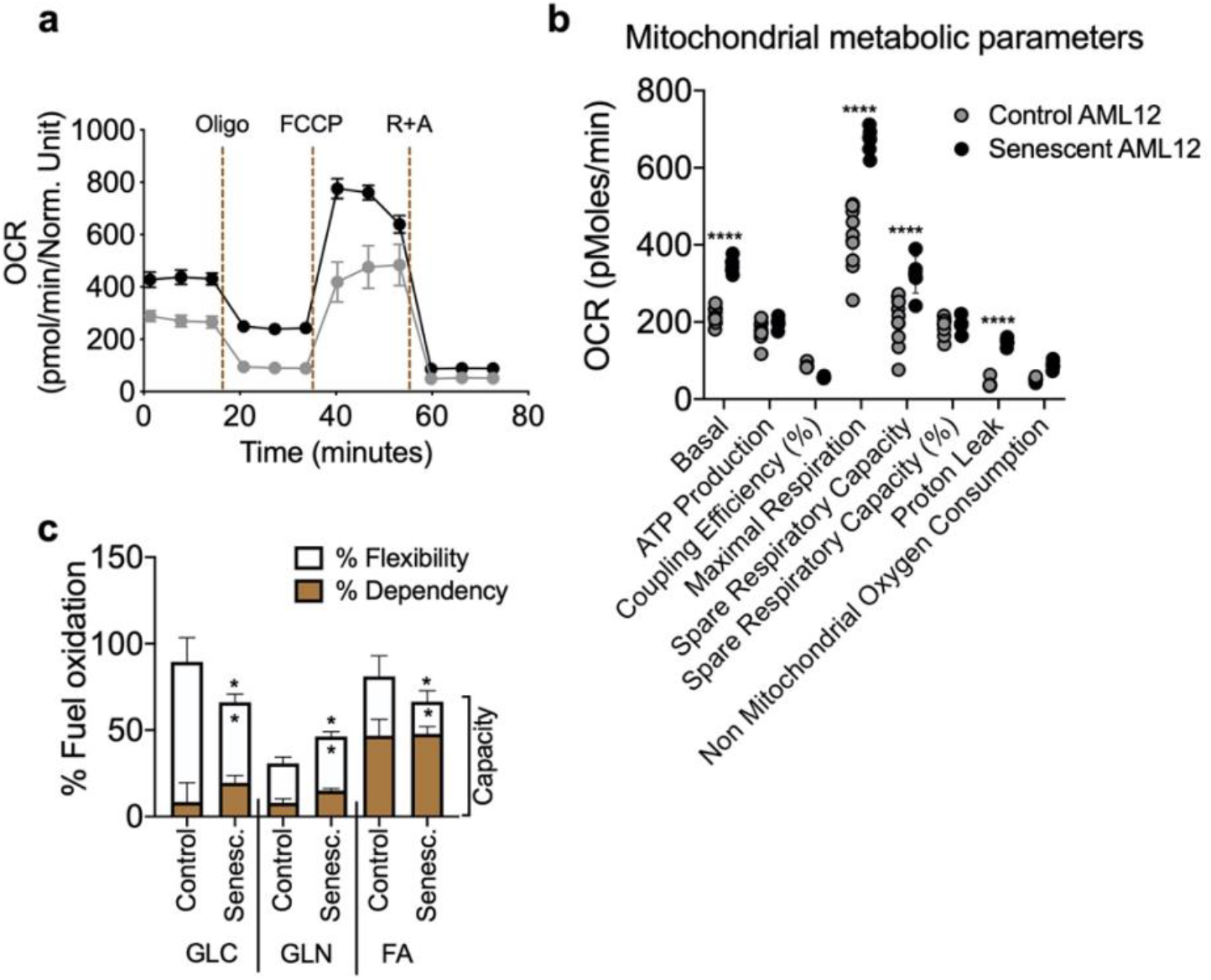
Seahorse extracellular flux analysis for mitochondrial metabolic parameters and fuel oxidation in senescent AML12 cells. Agilent Seahorse XF Mito Stress Test kit was used with Agilent Seahorse XFe96 Extracellular Flux Analyzer to analyse control and senescent AML12 cell’s mitochondrial metabolic potential (**a** and **b**). (**c**) Agilent Seahorse XF Mito Fuel Flex kit was used with Agilent Seahorse XFe96 Extracellular Flux Analyzer to analyse control and senescent AML12 cell’s mitochondrial fuel (glucose, glutamine and fatty acids) oxidation. All the parameters presented in the panel b and c were calculated using Seahorse Wave Desktop software. Statistical differences were calculated significant as *p<0.05 and ****p<0.0001.

### Molecular phenotyping of energy sensing molecules in senescent AML12 cells

We next analysed the activation (via phosphorylation) of several key energy-sensing proteins: AMPK, AKT, mTOR, p70S6K (mTOR target), as well as the formation of LC3B-II and p62 (autophagy-related proteins) in senescent AML12 cells by Western blotting. As a signature of senescent cells, we observed marked increases in the phosphorylation of AMPK, AKT, mTOR and p70S6K proteins in senescent AML12 cells (**Figure 5a,b**). This phosphorylation pattern was similar to the one observed in the livers of aged mice (**Figure 5c,d**). Furthermore, RNAseq pathway analysis also showed several insulin-related pathways along with prolong ERK1/2 and MAPK signaling pathways were upregulated (Supplementary figure 1b, Supplementary data-RNAseq Pathway analysis> Pathways Up), while negative regulation of mTOR signaling was downregulated (Supplementary figure 2a, Supplementary data-RNAseq Pathway analysis> Pathways Down). Additionally, we observed a marked increase in the expression of autophagy proteins, LC3B-II and P62, under basal conditions suggesting there was a late autophagic block in both the senescent AML12 cells and livers from aged mice. Furthermore, we confirmed autophagy block in senescent AML12 cells when a lysosome inhibitor bafilomycin A1 was used (**Figure 5e-f**). As evident by densitometric analysis that LC3B-II accumulation was declined in senescent AML12 cells. We also observed an accumulation of neutral lipids in senescent AML12 cells at basal conditions that was similar to the increased hepatic triglyceride content found in aged mice (Figure 5g-i). Consistently, RNAseq pathway analysis also showed several pathways regulating lipid handling, catabolism along with triglyceride catabolic pathways were downregulated (Supplementary figure 2a, Supplementary data-RNAseq Pathway analysis> Pathways Down).

**Figure 5.**
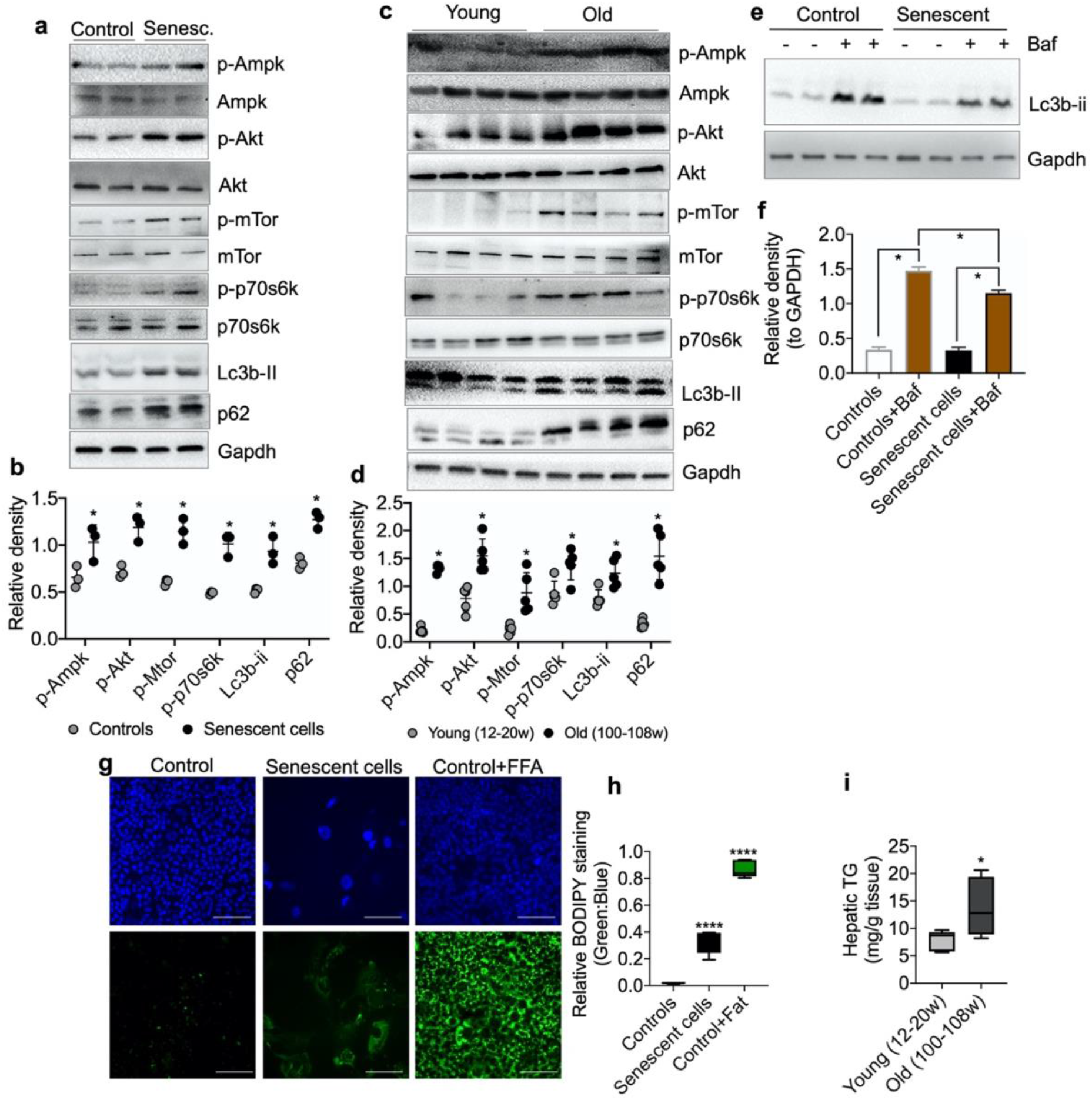
Molecular analysis of energy sensing pathways in senescent AML12 cells and liver tissues from young and old mice. (**a)** Western blot analysis of energy sensing and autophagic proteins including GAPDH in control and senescent AML12 cells. **(b**) Relative densitometric values of Western blots were calculated using ImageJ (NIH) software and normalized to GAPDH (n=3). (**c**) Western blot analysis of energy sensing and autophagic proteins including GAPDH in liver tissue from young and old mice. **(d**) Relative densitometric values of Western blots were calculated using ImageJ (NIH) software and normalized to GAPDH (n=5). (**e**) Western blot analysis of autophagy flux using lysosome inhibitor Bafilomycin A1 (Baf) in control and senescent AML12 cells under basal condition. (**f**) Relative densitometric values of Western blots were calculated using ImageJ (NIH) software and normalized to GAPDH (n=3). (**g**) Immunofluorescence analysis of neutral lipids accumulation (fat droplets) in control and senescent AML12 cells using BODIPY stain. Images were taken at 10x magnification. Control cells were treated overnight with 0.75 mM fatty acids Oleic Acid: Palmitic Acid (2:1) as positive control. Scale bars as 100 μm. **(h**) Relative BODIPY fluorescence was calculated over Hoechst 33342 fluorescence using ImageJ software (NIH). (**i**) Hepatic triglyceride measurement was performed in the liver tissues from young and old mice. Statistical differences were calculated significant as *p<0.05 and ****p<0.0001.

### Senescent AML12 cells create a pro-inflammatory environment and sensitize healthy AML12 cell towards pathological damage

Senescent cells secrete senescence-associated secretory proteins such as IL6 and IL1b, and are thought to be responsible for creating a pro-inflammatory environment for neighbouring cells during aging-associated diseases [1, 26, 27]. We found that senescent AML12 cells showed higher expression of the pro-inflammatory genes, *IL1b* and *IL6* (**Figure 6a**). Similarly, liver tissue from old mice showed significantly higher expressions of *IL1b* and *IL6* mRNA (**Figure 6b**). We next treated control AML12 cells with conditioned media collected from senescent AML12 cell cultures, and found that they not only increased the expression of inflammatory (*IL1b, IL6, CCL2*) and fibrosis (*IL11*) genes in normal AML12 cells but also increased the latter’s sensitivity to induction of inflammatory (*IL1b, IL6, CCL2*) and fibrosis (*IL11*) genes by saturated fatty acids (palmitate 0.5 mM for 24 h) in comparison to conditioned media prepared from control AML12 cell cultures (**Figure 6c**). Consistently, RNAseq pathway analysis revealed that IL1b production and IL1b secretory pathways along with extracellular matrix-related pathways were significantly upregulated (Supplementary figure 1d, Supplementary data-RNAseq Pathway analysis> Pathways Up). Furthermore, superoxide generation unfolded protein response (UPR), TNF-mediated signaling, and chemokine secretion pathways including urea cycle were also significantly upregulated (Supplementary figure 1a, Supplementary data-RNAseq Pathway analysis> Pathways Up). While, several glutathione metabolism regulating REDOX pathways along with nuclear receptors transcription pathways were significantly downregulated (Supplementary figure 2b-c, Supplementary data-RNAseq Pathway analysis> Pathways Down).

**Figure 6.**
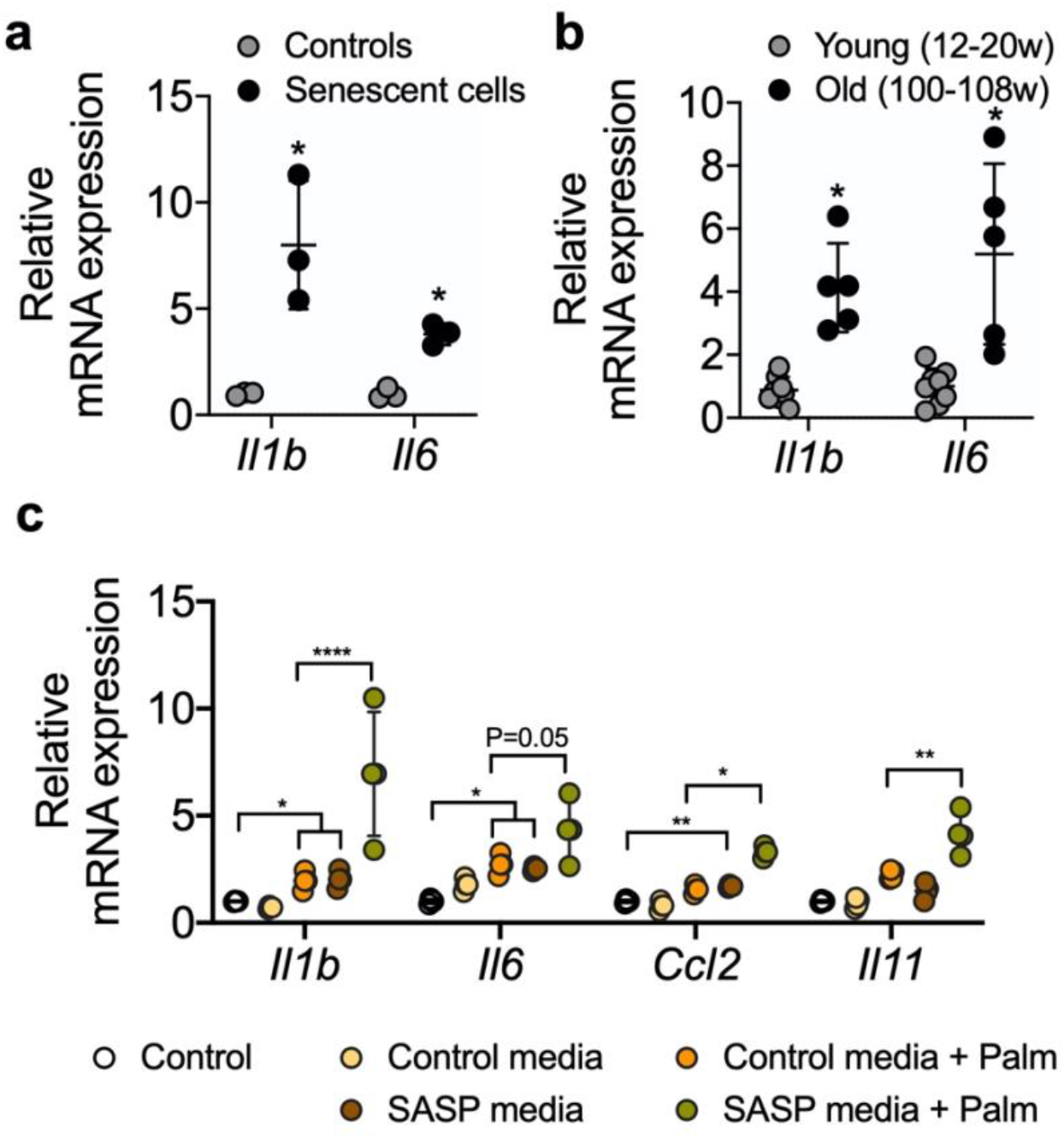
RT-qPCR analysis of SASP-related proinflammatory genes in senescent AML12 cells and liver tissues from young and old mice. RT-qPCR analysis of SASP-related proinflammatory genes in senescent AML12 cells (**a**) and liver tissues from young and old mice (**b**). (**c**) RT-qPCR analysis of inflammatory interleukins, chemokine CCL2 and fibrotic IL11 genes in control AML12 cells treated with conditional media from senescent AML12 cells (24 h) along with or without saturated fatty acids palmitate (0.5 mM for 24 h). Statistical differences were calculated significant as *p<0.05, **p<0.01, and ****p<0.0001.

## DISCUSSION

Both cellular senescence significantly increases in most tissues during aging [9, 12, 24]. Previous *in vitro* senescence models have utilized primary fibroblasts or hepatic carcinoma cells that were subjected to oxidative stress conditions or irradiation [17–23]. However, these models have limited application to the aging liver due to differences in their respective molecular and metabolic phenotypes. Since oxidative stress and mitochondrial dysfunction are considered to be critical for senescence and aging [6, 11, 24, 25], we subjected non-transformed AML12 hepatic cells to multiple exposures of sublethal H_2_O_2_ to induce premature cellular senescence to study hepatic aging *in vitro*. In particular, they showed increased expression of key senescence genes such as *Tp53, CDKN2A/p21*, and *CDKN1A*/*p16* as well as increased numbers of SA-βGal and γH2A.X positive cells that had condensed chromatin in larger nuclei (**Figures 1 and 2**).

Senescent cells are metabolically hyperactive and display a metabolic shift consisting of marked increases in glycolysis, mitochondrial activity, and mitochondrial damage due to proton leakage [8, 27, 28]. After AML12 cells were given multiple sublethal H_2_O_2_ doses, we found they increased their cellular bioenergetics (**Figure 3a,b**) and showed greater reliance upon glycolysis under nutrient stress conditions. Subsequent analysis of the glycolysis stress test confirmed senescent AML12 cells had greater glycolytic potential and reserve than control cells (**Figure 3c,d**). Senescent cells also have increased mitochondrial activity compared to control cells. However, this increased activity did not lead to more ATP production due to concomitant increases in basal uncoupling and proton leakage that occurred in senescent cells [1, 25]. These findings suggested that increased intracellular AMP:ATP concentration ratio in senescent cells led to the AMPK activation [1, 27, 29].

The senescent AML12 cells also had significantly higher basal respiration and maximum respiration than normal AML 12 cells (**Figure 4a,b**). However, these changes were not accompanied by increases in % spare respiratory capacity and ATP production. Notably, similar changes in mitochondrial function also have been reported to be present in livers from aged mice [1]. Previously, impairments in hepatic fatty acid oxidation and glucose intolerance have been described in the aged liver [1, 30]. In the senescent AML12 cells, mitochondrial fuel oxidation analysis also showed significantly lower fuel flexibility and less total capacity for glucose and fatty acid oxidation (**Figure 4c**). Taken together, our findings show that there is altered metabolism and mitochondrial function in senescent AML12 cells that resemble some of the changes previously reported in the aged liver [1, 8, 27, 28, 30].

Interestingly, senescent AML12 cells also showed a compensatory increase in mitochondrial glutamine oxidation (**Figure 4c**). Glutamine activates hepatic metabolic pathways involving PEPCK and the p70S6K. Phosphorylation of the latter enzymes leads to inhibition of autophagic proteolysis and induction of cell swelling [31, 32]. Glutamine oxidation also facilitates mTOR activation by leucine, an activator of glutaminolysis, to regulate cell growth and autophagy [33]. In this connection, we also observed decreased autophagy in senescent AML12 cells (**Figure 5a,b**). It is likely that the activated mTOR signaling and increased p70S6K phosphorylation led to the inhibition of autophagy. Moreover, this decreases in autophagy fatty acid oxidation likely contributed to the fuel switching since we previously showed that both lipophagy and mitophagy are critical for β-oxidation of fatty acids in the liver [34–36]. Supporting this notion, we saw increased fat accumulation in the senescent cells (**Figure 5e-g**). Our findings also were consistent with the observation that autophagy impairment associated with lysosomal and mitochondrial dysfunction is an important characteristic of oxidative stress-induced senescence [37]. On the other hand, our findings are in contrast to several studies that showed little or no change in autophagy during senescence [28, 38]. These differences suggest that may be cell-specific changes in autophagy during senescence. Nonetheless, it appears that in the liver, autophagy likely decreases with aging and may be involved in many age-related diseases such as NAFLD and Type 2 DM [1, 3, 11, 12, 39]. In this connection, we found that both senescent AML12 cells and livers from aged mice showed late autophagy block associated with increased intracellular fat content (**Figure 5a-g**).

AMPK is a glucose and cellular energy sensor that is activated during states of energy depletion. In the liver, it is able to sense increased AMP:ATP and ADP:ATP concentration ratios, and activates compensatory responses such as increasing fatty acid oxidation, mitochondrial biogenesis, and glucose uptake, as well as inhibiting fatty acid synthesis [40, 41]. AMPK activation also induces cell cycle arrest and senescence by directly phosphorylating p53 at multiple sites [42]. In contrast to AMPK, which senses energy depletion and activates catabolic pathways, mTOR senses the fed state and activates anabolic pathways during cell growth and proliferation. Thus, AMPK and mTOR often have opposing roles in the metabolically active tissues of mammals [43]. However, the reciprocal actions of AMPK and mTOR that maintain metabolic homeostasis becomes impaired during senescence, and leads to concurrent activation of both mTOR and AMPK. This dysregulation also is found in the aged liver [1, 27, 28]. Similarly, we found that both AMPK and mTOR were activated in senescent AML12 cells (**Figure 5**), suggesting that energy sensing became dysregulated due to impaired oxidative phosphorylation, ATP production, (**Figure 4a,b**), and a compensatory increase in anaerobic glycolysis (**Figure 3c,d**).

Senescent cells employ an energy-dependent process to secrete senescence associated secretory proteins comprised of high levels of inflammatory cytokines, immune modulators, growth factors, and proteases [7, 8, 28, 44, 45]. IL6 and IL1β are major secretory interleukins that are present in senescence associated secretory phenotype (SASP) [1, 3, 8, 46]. In this connection, we observed that glycolysis increased in senescent AML12 cells (**Figure 3c,d**), and was accompanied by increased expression of IL6 and IL1β mRNA. Similar findings also were observed in livers from aged mice (**Figure 6a,b**). Interestingly, when we incubated healthy AML12 cells with conditional media collected from senescent AML12 cells, they became more sensitive to the inflammatory response caused by the saturated fatty acid, palmitate (**Figure 6c**). Our findings showed that senescence associated secretory proteins play an important role in generating a pro-inflammatory environment during aging and age-related metabolic diseases, and were consistent with earlier reports on SASP [1, 3, 8, 46].

In conclusion, we generated senescent AML12 hepatic cells that faithfully recapitulate many of the features of senescence found in livers of aged mice. In particular, the senescent AML12 cells expressed many of the key cellular markers of senescence, and exhibited altered mitochondrial metabolism and molecular signaling that were similar to those found in the livers of aged mice. Most interestingly, they showed metabolic fuel switching from fatty acid utilization in normal AML12 cells to glycolysis and glutamine utilization. The increase in mTOR signaling, perhaps due to increased glycolysis, led to a decrease in autophagy which then decreased β-oxidation of fatty acids. This process further increased glycolysis and increased glutamate oxidation by the senescent AML12 cells. Furthermore, the senescent AML12 cells produced a pro-inflammatory environment that rendered neighbouring hepatic cells more susceptible to saturated fatty acid-induced toxicity. The latter effect may occur in NAFLD during the progression from hepatosteatosis to non-alcoholic steatohepatitis (NASH). In summary, senescent AML 12 cells share many of the molecular, cell signaling, and metabolic characteristics found in the aged liver, and will be useful tool for further mechanistic studies on aging in the liver.

## METHODS

### Senescence induction in normal mouse AML12 cells and analysis of cell replication

The AML12 (alpha mouse liver 12) cells were purchased from ATCC, USA (ATCC® CRL-2254™) and maintained as described in the standard protocol (https://www.atcc.org/Products/All/CRL-2254.aspx#culturemethod). 4×10_6_ AML12 cells were seeded in a T175 cell culture flask for senescence induction. 30% (9.77 M) hydrogen peroxide (H_2_O_2_) was used for senescence induction. At day 1, cells were treated with 1 mM H_2_O_2_ for 1 h in serum-free medium followed by incubation in complete DMEM:F12 medium (containing 10% FBS, 1x ITS, 100 nM dexamethasone, and 1x penicillin and streptomycin) for recovery for 23 h. from day 2 to day 6, 750 μM H_2_O_2_ for 1 h in serum-free medium followed by 23 h recovery was used. Morphological changes should be visualized from day 3 of H_2_O_2_ treatment (**figure 1b**). At day 7, 0.3×10_6_ control and senescent AML12 cells were seeded in each well of 6-well plate for cell replication analysis and counted (using Countess™ II Automated Cell Counter, ThermoFisher) at every other day for three days as shown in **figure 1c**.

### Senescence activated secretory phenotype (SASP)-induced palmitate toxicity

AML12 senescence was induced as described above and after day 7 AML12 complete media was replaced with basal media (DMEM:F12) containing only 1x penicillin and streptomycin for 24 h. The next day, this conditional media was collected from control and senescent AML12 cells and put on new control AML12 cells seeded in other 6-well plates for 24 h. Palmitate was used at 0.5 mM concentration in 0.5% BSA supplemented (in a pathological FA:BSA ratio 6:1) in control media as well as senescent conditional media was used to represent pathological fatty acids concentration in serum [47]. Equivalent BSA was used in appropriate controls.

### Animal experiments

12-20 weeks age male C57BL6/J mice (consider young), whereas 100-108 weeks age male C57BL6/J mice (consider old) were purchased from Jacksons Laboratory USA (Stock: 000664) and used in this study. Animals were housed in hanging polycarbonate cages under a 12 h light/12 h dark cycle at 23 °C with food and water available *ad libitum*. All cages contained shelters and nesting material. 6 h fasted mice euthanized and liver was collected and snap frozen in liquid nitrogen for subsequent analysis. All mice were maintained according to the Guide for the Care and Use of Laboratory Animals (National Institutes of Health publication 1.0.0; revised 2011), and experiments were approved by SingHealth Institutional Animal Care and Use Committee.

### mRNA expression analysis by reverse transcription-quantitative PCR

Total RNA isolation from cultured cells and liver tissue was performed using InviTrap Spin Universal RNA kit (Stratec Biomedical, Birkenfeld, Germany), and RT-qPCR was performed as described previously [48] using QuantiTect SYBR Green PCR kit and KiCqStart SYBR Green optimized primers from Sigma-Aldrich (KSPQ12012).

### Protein extraction and expression analysis by Western blotting

Cultured cells or 50 mg of liver tissues were lysed using CelLytic M mammalian cell lysis/extraction reagent (C2978, Sigma-Aldrich). An aliquot was removed, and protein concentrations were measured using the BCA kit (Bio-Rad). Western blotting was performed using a standard protocol, as described previously [49]. Primary antibodies at dilution 1:500 for phospho-AMPK-alpha (T172) (CST: 2535S), AMPK-alpha (D5A2) (CST: 5831S), phospho-AKT (S473) (CST: 4058S), AKT (CST: 9272S), phospho-mTOR (S2448) (CST: 5536S), mTOR (7C10) (CST: 2983S), phospho-p70S6K (S371) (CST: 9206S), p70S6K (CST: 9202s) 1:5000 for LC3B-II (CST: 2775S), GAPDH (CST: 2118L), and 1:1000 for p62 (CST: 5114S) were used. Western blot images were captured on the Gel-Doc system (Bio-Rad), and densitometry analysis was performed using ImageJ software (National Institutes of Health). The integrated density of target protein was normalized with the GAPDH; the mean was plotted in graphs.

### Immunofluorescence analysis for γH2A.X

Cells were cultured in 4-well chambered slides, fixed in 4% paraformaldehyde and incubated in 1:200 diluted γH2A.X antibody (CST: 9718) overnight at 4 _o_C after blocking as described previously [36]. 1:200 Alexa Fluor 488 (Molecular Probes, ThermoFisher) was used to collect signals. Cells were counter-stained with 5 μM Hoechst 33342 (Abcam: ab228551) for 5 min and mounted using VECTASHIELD_®_ Antifade Mounting Medium H-1000 (Vector Laboratories), and visualized under 40x magnification using Zeiss LSM confocal microscope. Five images were clicked randomly using ZEN 2 (blue edition) software and quantification of green fluorescence was normalized with blue using ImageJ software (National Institutes of Health) and plotted as a graph.

### Senescent associated β-Gal staining in AML12 cells

Cellular Senescence Assay kit (Merck Millipore: KAA002) was used to detect senescent AML12 by SA β-Gal staining as per the manufacturer’s protocol. X-Gal staining was performed overnight at 37 _o_C and crystals were visualized under the microscope at 10x magnification. Five images were captured randomly using Olympus inverted microscope and cells were counted for SA β-Gal_+_ Cells, and shown as percent positive cells.

### Seahorse extracellular flux analysis for bioenergetic phenotyping, mitostress test, glycolytic stress test and mitochondrial fuel oxidation analysis

Seahorse extracellular flux analyser XFe96 (Agilent) was used for bioenergetic phenotyping, mitostress test, glycolytic stress test and mitochondrial fuel oxidation analysis, and Seahorse Wave Desktop software was used for report generation and data analysis, and GraphPad PRISM 8 was used for statistical analysis and data presentation.

Bioenergetic phenotyping: Agilent Seahorse XF Cell Energy Phenotype Test kit was used with Agilent Seahorse XFe96 Extracellular Flux Analyzer that rapidly measures mitochondrial respiration and glycolysis under baseline and stressed conditions, to reveal the three key parameters of cell energy metabolism: Baseline Phenotype, Stressed Phenotype, and Metabolic Potential (https://www.agilent.com/cs/library/usermanuals/public/XF_Cell_Energy_Phenotype_Test_Kit_User_Guide.pdf). Simultaneous acute injections of mitochondrial inhibitors oligomycin and FCCP reveals live cell’s metabolic potential under stress. Oligomycin (1 μM) inhibits ATP production by the mitochondria, and causes a compensatory increase in the rate of glycolysis as the cells attempt to meet their energy demands via the glycolytic pathway. Whereas, FCCP (1 μM) depolarizes the mitochondrial membrane, and drives oxygen consumption rates higher as the mitochondria attempt to restore the mitochondrial membrane potential.

Glycolysis stress test: Agilent Seahorse XF Glycolysis Stress Test Kit was used as per the standard protocol by Agilent Seahorse (https://www.agilent.com/en-us/agilent404?s=www.agilent.com/cs/library/usermanuals/public/XF_Glycolysis_Stress_Test_Kit_User_Guide.pdf). Glucose conversion to lactate (Glycolysis) results in net production and extrusion of protons into the extracellular medium (acidification). As glycolysis occurs, the resulting acidification of the medium surrounding the cells is measured directly by the analyzer and reported as the Extracellular Acidification Rate (ECAR).

To understand the complete glycolytic potential of the cell, ECAR of cells was measured (three times) before and after sequential injections of three different compounds, glucose (10 mM), oligomycin (1 μM) and 2-Deoxy-D-glucose (2-DG; 100 mM) sequentially.

Mito stress test: Agilent Seahorse XF Mito Stress Test Kit was used as per the standard protocol by Agilent Seahorse (https://www.agilent.com/cs/library/usermanuals/public/XF_Cell_Mito_Stress_Test_Kit_User_Guide.pdf). This test measures key parameters of mitochondrial function by directly measuring the oxygen consumption rate (OCR) of cells. Oligomycin (1 μM; inhibits ATP synthase) was injected first following basal OCR measurements. The second injection of FCCP (1 μM; a uncoupling agent that collapses the proton gradient and disrupts the mitochondrial membrane potential) followed by the third injection which was a mixture of rotenone (1 μM; a complex I inhibitor) and antimycin A (1 μM; a complex III inhibitor).

Mito fuel flex test: Agilent Seahorse XF Mito Fuel Flex Test Kit was used as per the standard protocol by Agilent Seahorse (https://www.agilent.com/cs/library/usermanuals/public/XF_Mito_Fuel_Flex_Test_Kit_User_Guide%20old.pdf). This test measures the dependency, capacity, and flexibility of cells to oxidize three mitochondrial fuels in real-time in living cells: Glucose (pyruvate), Glutamine (glutamate) and Long-chain fatty acids. This test determines the rate of oxidation of each of these fuels by measuring OCR of cells in the presence or absence of fuel pathway inhibitors. UK5099 (2 μM; glucose oxidation pathway inhibitor) that blocks the mitochondrial pyruvate carrier (MPC); BPTES (3 μM; glutamine oxidation pathway inhibitor) that allosterically inhibits glutaminase (GLS1); and Etomoxir (4 μM; long chain fatty acid oxidation inhibitor) that inhibits carnitine palmitoyl-transferase 1A (CPT1A), a critical enzyme of mitochondrial beta-oxidation.

The data later represented as fuel oxidation dependency, flexibility and total capacity. ‘Dependency’ indicates the fuel oxidation at basal in control or senescence cells. Inhibiting the two alternative pathways followed by the pathway of interest enables the calculation of cells’ mitochondrial ‘capacity’ to meet energy demand. ‘Flexibility’ (that is calculated by subtracting the dependency from the capacity for the target fuel oxidation pathway) indicates the cells’ mitochondria can compensate for the inhibited pathway by using other pathways to fuel mitochondrial respiration. The presence of dependency and the absence of flexibility demonstrates that the mitochondria require that fuel pathway to maintain basal OCR.

### Immunofluorescence imaging of intracellular lipids using BODIPY™ 493/503

Control and senescent AML12 cells were seeded in 24-well plate at 7_th_ day of the protocol, and cultured for 24 h. Cells were rinsed with 1x PBS and stained with BODIPY™ 493/503 (D3922; Molecular Probes, ThermoFisher Scientific) at 1:1000 dilution for 15 min. Cells were then rinsed with 1xPBS containing Hoechst 33342 (Sigma) to counter stain nucleus for 5 min. After rinse, cells were kept in 1x HBSS for imaging. Leica fluorescent microscope was used at 10x magnification for visualization and LAS X imaging software was used for image capture.

### Liver triglycerides measurements in young and old mice

Liver triglyceride was measured using triglyceride colorimetric assay kit (Cayman Chemical) as per the manufacturer’s protocol.

### RNAseq and pathway analysis

The RNAseq dataset is submitted as Series record GSE151806 on Gene Expression Omnibus, NCBI. Detailed method on RNAseq analysis is provided as supplementary methods. Significant (p<0.05) deferentially expressed genes with threshold more than 1.5 log2fold change or less than −1.5 log2fold change were analysed for upregulated or downregulated pathway using EnrichR online pathway analysis platform (https://amp.pharm.mssm.edu/Enrichr/) [50, 51]. Pathways were captured from KEGG 2019 Human, Reactome 2016, as well as Gene Ontology (GO) biological process and molecular functions databases.

## Supporting information

liked to Supplementary figure 1 and 2.

## Statistical methods

The data was calculated and presented as Mean±SD. The parametric unpaired t-test was use to compute significance between two groups, whereas Two-way ANOVA followed by Tukey’s multiple comparisons test was used to compute significance between more than two groups. GraphPad PRISM 8 was used for statistical analysis and data representation.

## ACKNOWLEDGMENTS

The authors would like to thank Ms. Jia Pei and Mr. Kiraely Adam Wong Zongren for their assistance and help during this study.

## AUTHOR CONTRIBUTIONS

Conceived and designed the experiments: BKS MT. Performed the experiments: BKS MT RS KT. Analyzed the data: BKS MT JZ. Contributed reagents/materials/analysis tools: PMY BKS MT JZ. Wrote the manuscript: BKS MT PMY.

## SUPPLEMENTARY FIGURE LEGENDS

**Figure 1.**
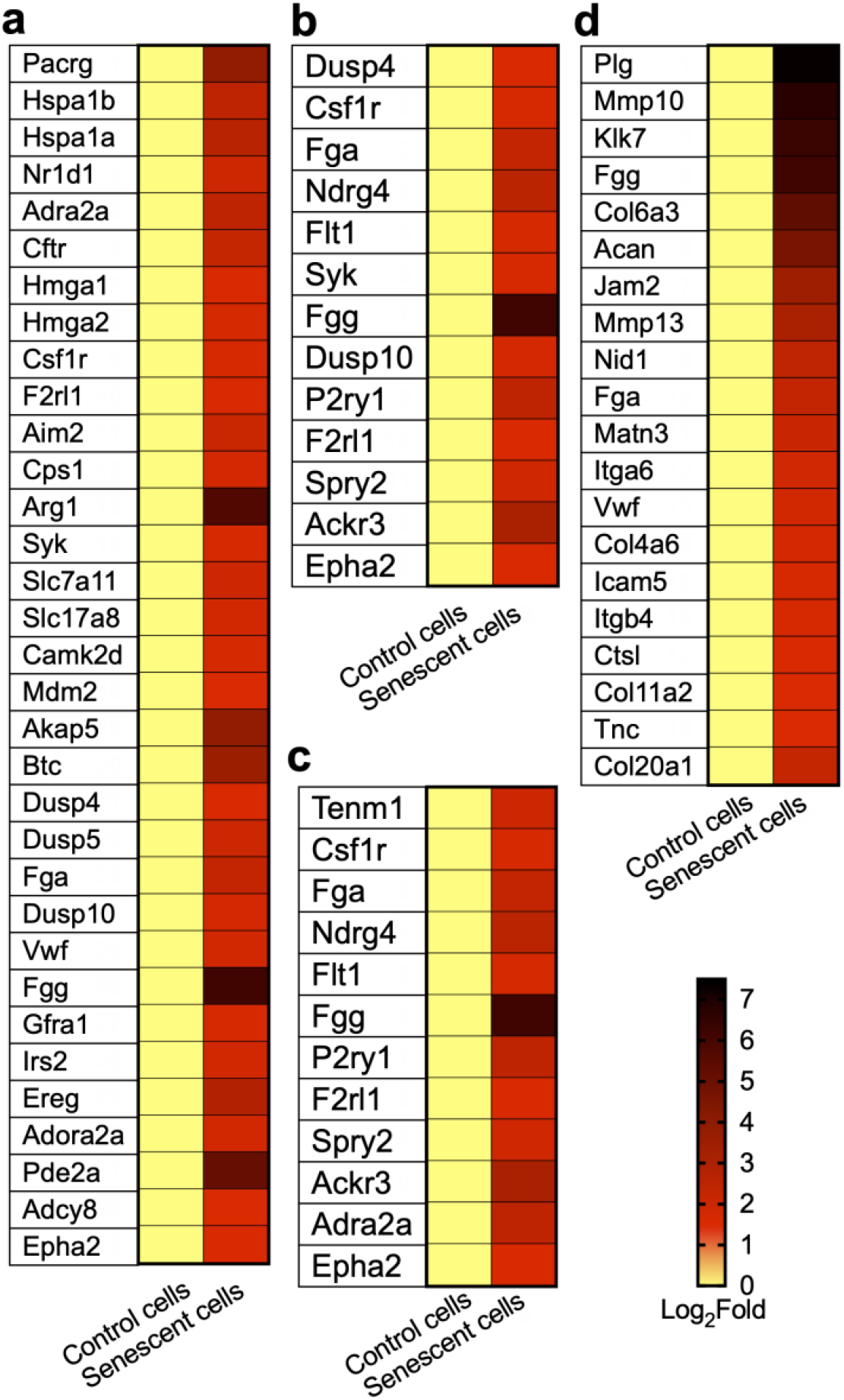
RNAseq analysis of control and senescent AML12 cells for upregulated pathways. (**a**) Heat map showing genes regulating pathways: cAMP metabolic process (GO:0046058), positive regulation of tumor necrosis factor-mediated signaling pathway (GO:1903265), cellular response to unfolded protein (GO:0034620), regulation of insulin secretion involved in cellular response to glucose stimulus (GO:0061178), positive regulation of cellular senescence (GO:2000774), positive regulation of cell aging (GO:0090343), regulation of chemokine secretion (GO:0090196), interleukin-1 beta secretion (GO:0050702), interleukin-1 beta production (GO:0032611), bile acid biosynthetic process (GO:0006699), urea cycle (GO:0000050), regulation of superoxide anion generation (GO:0032928), L-glutamate transmembrane transporter activity (GO:0005313), Glutamate Binding, Activation of AMPA Receptors and Synaptic Plasticity Homo sapiens R-HSA-399721, Insulin receptor signalling cascade Homo sapiens R-HSA-74751, Signaling by Type 1 Insulin-like Growth Factor 1 Receptor (IGF1R) Homo sapiens R-HSA-2404192, IRS-mediated signalling Homo sapiens R-HSA-112399, and Signaling by Insulin receptor Homo sapiens R-HSA-74752. (**b**) Heat map showing genes regulating pathways: regulation of ERK1 and ERK2 cascade (GO:0070372), and Prolonged ERK activation events Homo sapiens R-HSA-169893. (**c**) Heat map showing genes regulating pathway: positive regulation of MAPK cascade (GO:0043410).

**Figure 2.**
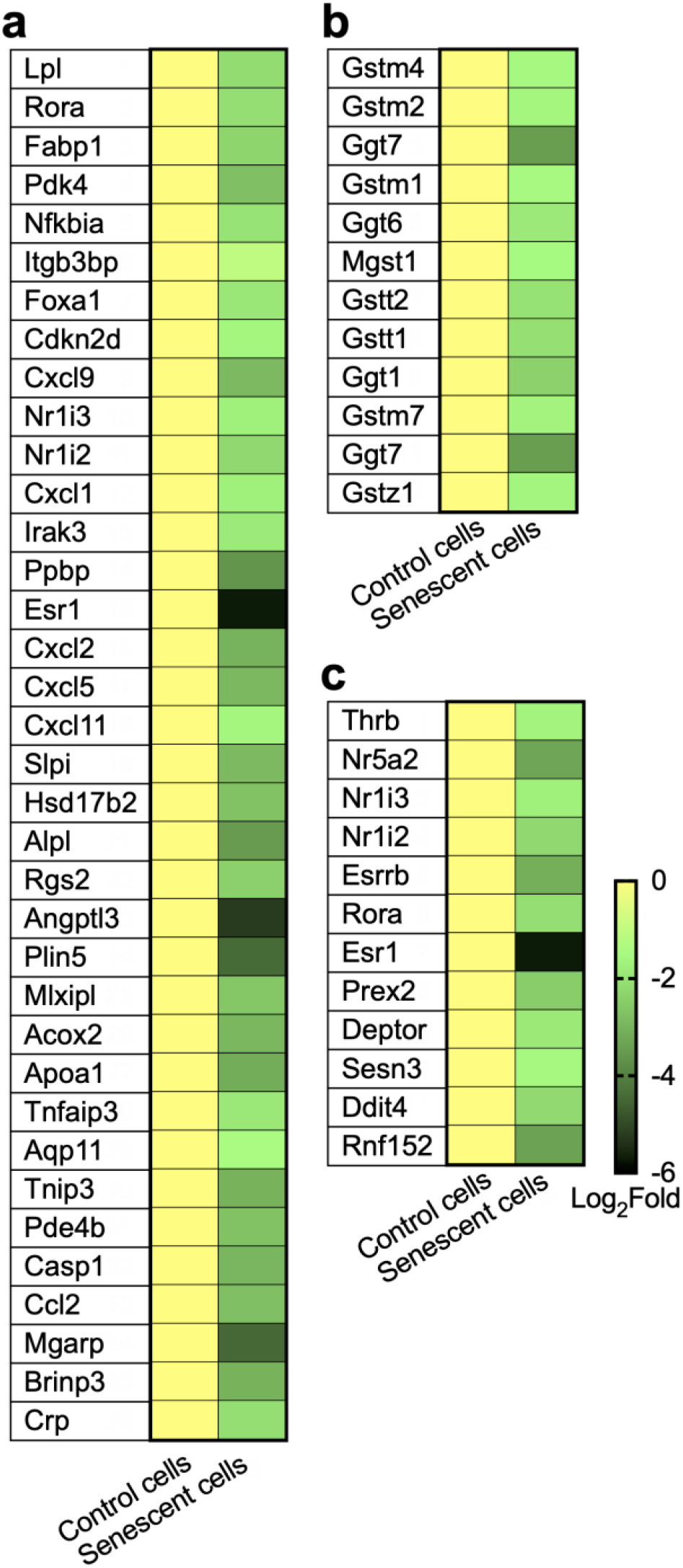
RNAseq analysis of control and senescent AML12 cells for downregulated pathways. (**a**) Heat map showing genes regulating pathways: response to lipid (GO:0033993), negative regulation of lipase activity (GO:0060192), lipid homeostasis (GO:0055088), triglyceride homeostasis (GO:0070328), regulation of cholesterol homeostasis (GO:2000188), triglyceride catabolic process (GO:0019433), cellular response to lipid (GO:0071396), negative regulation of lipid storage (GO:0010888), and regulation of lipid biosynthetic process (GO:0046890). (**b**) Heat map showing genes regulating pathways: Glutathione metabolism, Glutathione conjugation Homo sapiens R-HSA-156590, Glutathione synthesis and recycling Homo sapiens R-HSA-174403, glutathione derivative biosynthetic process (GO:1901687), and glutathione metabolic process (GO:0006749). (**c**) Heat map showing genes regulating pathways: Nuclear Receptor transcription pathway Homo sapiens R-HSA-383280, and negative regulation of TOR signaling (GO:0032007).

